# A Four-Step Enzymatic Cascade for Efficient Production of L-Phenylglycine from Biobased L-Phenylalanine

**DOI:** 10.1101/2022.01.14.476296

**Authors:** Yuling Zhu, Jifeng Yuan

**Affiliations:** State Key Laboratory of Cellular Stress Biology, School of Life Sciences, Xiamen University, Fujian 361102, PR China

**Keywords:** L-phenylglycine, L-amino acid deaminase, multi-enzyme cascade, leucine dehydrogenase, whole-cell biocatalyst.

## Abstract

Enantiopure amino acids are of particular interest in the agrochemical and pharmaceutical industries. Here, we reported a multi-enzyme cascade for efficient production of L-phenylglycine (L-Phg) from biobased L-phenylalanine (L-Phe). We first attempted to engineer *Escherichia coli* for expressing L-amino acid deaminase (LAAD) from *Proteus mirabilis*, hydroxymandelate synthase (HmaS) from *Amycolatopsis orientalis*, (*S*)-mandelate dehydrogenase (SMDH) from *Pseudomonas putida*, the endogenous aminotransferase (AT) encoded by *ilvE* and L-glutamate dehydrogenase (GluDH) from *E. coli*. However, 10 mM L-Phe only afforded the synthesis of 7.21 ± 0.15 mM L-Phg. The accumulation of benzoylformic acid suggested that the transamination step might be rate-limiting. We next used leucine dehydrogenase (LeuDH) from *Bacillus cereus* to bypass the use of L-glutamate as amine donor, and 40 mM L-Phe gave 39.97 ± 3.84 mM (6.04 ± 0.58 g/L) L-Phg, reaching 99.9% conversion. In summary, this work demonstrated a concise four-step enzymatic cascade for the L-Phg synthesis from biobased L-Phe, with a potential for future industrial applications.

**Graphical abstract:** a concise four-step enzymatic cascade for the L-phenylglycine synthesis from biobased L-phenylalanine was devised. 40 mM L-phenylalanine afforded the synthesis of 39.97 ± 3.84 mM (6.04 ± 0.58 g/L) L-phenylglycine, reaching 99.9% conversion.

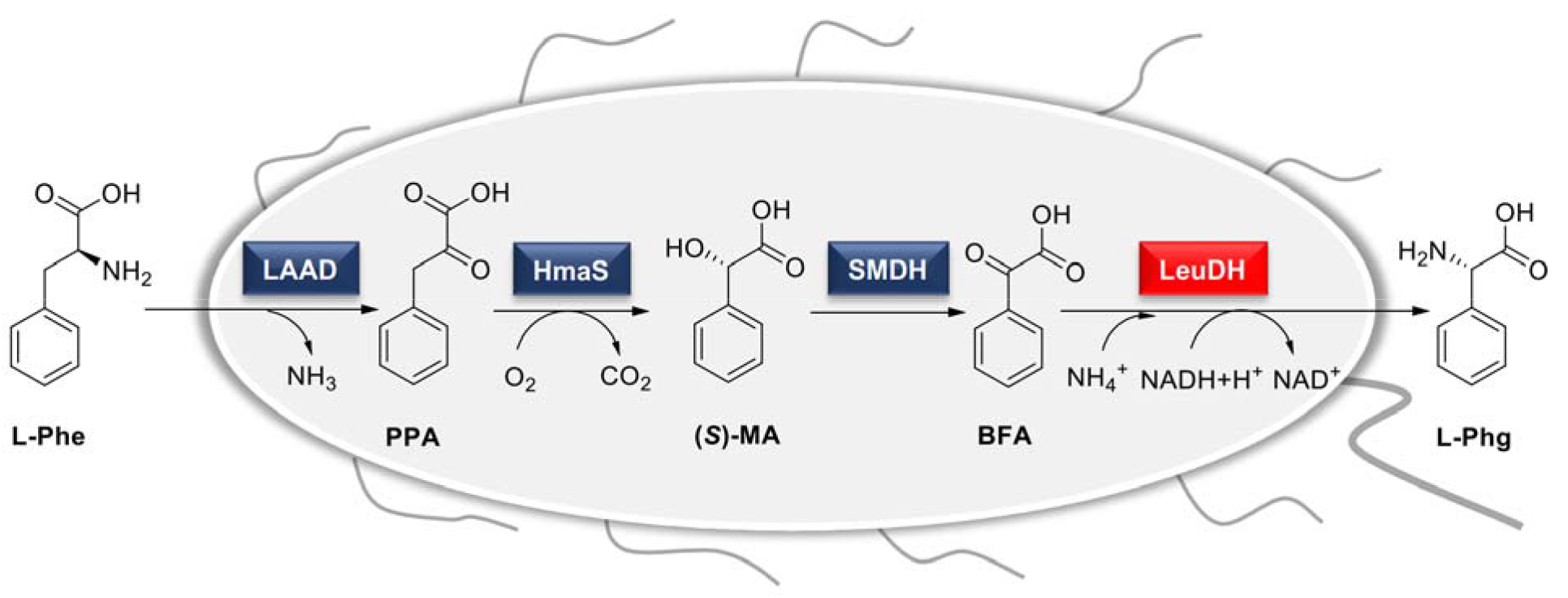

---

Enantiopure amino acids are of particular interest in the agrochemical and pharmaceutical industries ^[1]^. For example, L-phenylglycine (L-Phg) can be used for the synthesis of β-lactam antibiotics such as penicillin ^[2]^, streptogramin B ^[3]^, pristinamycin I ^[4]^ and virginiamycin S ^[5]^. A number of chemical synthesis routes have been developed for the production of enantiopure amino acid ^[6]^. For instance, the Strecker reaction is a common method for the industrial production of α-amino acids ^[7]^. However, chemical approaches typically use environmentally harmful toxic reagents and organic solvents, and the enantioselectivity of the product is relatively low, which are undesirable for sustainable production of enantiopure amino acids ^[8]^.

In comparison to chemical approaches, biocatalytic processes are usually carried out under less harsh conditions such as ambient temperature and atmospheric pressure ^[9]^. In addition, biocatalytic processes are considered as a suitable alternative for chiral chemical productions because of the excellent enantioselectivity ^[10]^. Since amino acids such as L-phenylalanine (L-Phe) can be obtained by fermentation or from protein waste hydrolysates in large amounts and at a low cost ^[11]^, they are considered to be cheap and renewable feedstocks for biomanufacturing applications. For instance, it was reported that the production of L-Phg from biobased L-Phe was achieved in the recombinant *Escherichia coli* containing eight reaction steps with 12 genes ^[12]^. Under the optimal condition, 40 mM L-Phe was converted to 34 mM (5.1 g/L) L-Phg after 24 h via the whole-cell biotransformation, reaching ~85% conversion. In addition, *de novo* synthesis of L-Phg ^[14]^ and *in vitro* enzyme catalysis ^[15]^ have also been reported for the L-Phg production, the L-Phg yields were relatively low (51.6 mg/g DCW and 91.4 mg/L, respectively).

In this study, we sought to develop a more concise multi-enzyme cascade for synthesizing L-Phg from biobased L-Phe. In particular, our system comprises L-amino acid deaminase (LAAD, Uniprot ID: B2ZHY0) from *Proteus mirabilis* ^[13]^, hydroxymandelate synthase (HmaS, Uniprot ID: O52791) from *Amycolatopsis orientalis* ^[14]^, (*S*)-mandelate dehydrogenase (SMDH, Uniprot ID: P20932) from *Pseudomonas putida*^[15]^ and leucine dehydrogenase (LeuDH, Uniprot ID: P0A393) from *Bacillus cereus* ^[16]^. The four-step enzymatic cascade could efficiently convert 40 mM L-Phe to 39.97 ± 3.84 mM (6.04 ± 0.58 g/L) L-Phg after 12 h, with a conversion rate >99.9%. To the best of our knowledge, our work represents one of the most efficient biocatalytic routes for L-Phg synthesis from L-Phe ^[12]^.

Recently, our group have demonstrated that LAAD from *P. mirabilis* and HmaS from *A. orientalis* together with SMDH from *P. putida* could effectively synthesize benzyl alcohol and benzylamine from L-Phe ^[17]^. To enable L-Phg production from benzoylformic acid (BFA), we chose aminotransferase (AT, Uniprot ID: P0AB80) encoded by *ilvE* and L-glutamate dehydrogenase (GluDH, Uniprot ID: P00370) from *E. coli* ^[12], [18]^. In brief, the multi-enzyme cascade comprises LAAD from *P. mirabilis*, HmaS from *A. orientalis*, SMDH from *P. putida* and AT/GluDH from *E. coli* (Scheme 1). As depicted in Figure 1a, the multi-enzyme cascade was recast into three modules: LAAD together with HmaS to convert L-Phe into (*S*)-mandelic acid; SMDH and AT to convert (*S*)-mandelic acid into L-Phg; an additional plasmid expressing GluDH from *E. coli* ^[12], [18]^ to improve the L-glutamate regeneration. When the recombinant *E. coli* strain Ec-Phg1.0 harboring three plasmids (pET-LAAD- HmaS, pRSF-SMDH-AT and pACYC-GluDH) was analyzed by the SDS-PAGE, we could clearly observe the bands corresponding to HmaS, SMDH, AT and GluDH (Figure 1b). However, LAAD was not observed from the SDS-PAGE result, probably because LAAD is a membrane-associated protein which is less abundant.

**Scheme 1.**
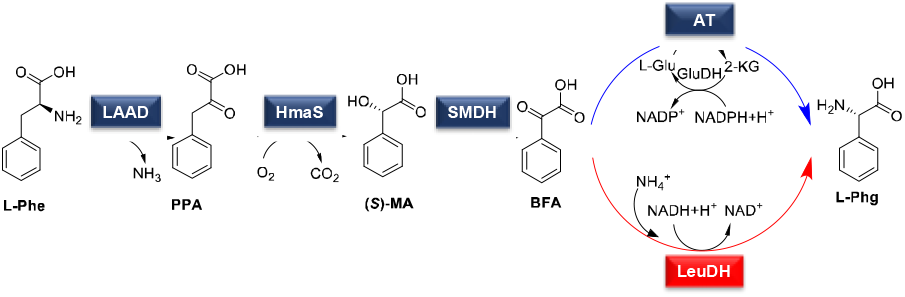
Schematic diagram of multi-enzyme cascade toward L-Phg synthesis from biobased L-Phe. LAAD, L-amino acid deaminase from *P. mirabilis* (Uniprot ID: B2ZHY0); HmaS, hydroxymandelate synthase from *A. orientalis* (Uniprot ID: O52791); SMDH, (*S*)-mandelate dehydrogenase from *P. putida* (Uniprot ID: P20932); AT, aminotransferase encoded by *ilvE* from *E. coli* (Uniprot ID: P0AB80); GluDH, L-glutamate dehydrogenase from *E. coli* (Uniprot ID: P00370); LeuDH, leucine dehydrogenase from *B. cereus* (Uniprot ID: P0A393). L-Phe, L-phenylalanine; PPA, phenylpyruvate; (*S*)-MA, (*S*)-mandelic acid; BFA, benzoylformic acid; L-Phg, L-phenylglycine; 2-KG, 2-ketoglutarate; L-Glu, L-glutamate.

**Figure 1.**
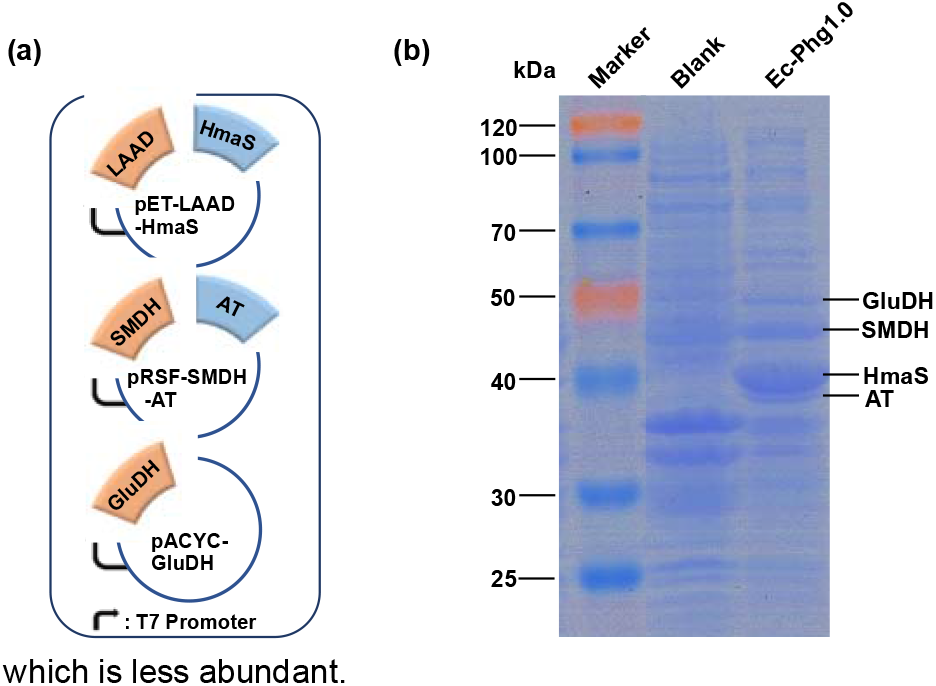
Plasmid design of aminotransferase (AT)-dependent route and SDS-PAGE analysis of protein expression. (a) The recombinant *E. coli* strain Ec-Phg1.0 containing three plasmids (pET-LAAD-HmaS, pRSF-SMDH-AT and pACYC-GluDH). (b) SDS-PAGE analysis of the recombinant *E. coli* strain Ec-Phg1.0. Blank: *E. coli* BL21 (DE3) harboring empty plasmids.

In the early experiments, we compared the effect of different pH conditions on L-Phg production, and we found that the ideal pH for L-Phg synthesis was around 8.0 (Figure 2a). As shown in Figure 2b and Table S1, when 10 mM L-Phe was fed to the recombinant *E. coli* Ec-Phg1.0 expressing five enzymes, 7.21 ± 0.15 mM L-Phg was obtained after 48 h, which corresponds to ~72.1% conversion. From the HPLC result, we found a substantial amount of BFA was accumulated during the biocatalytic process (Figure 2c).

**Figure 2.**
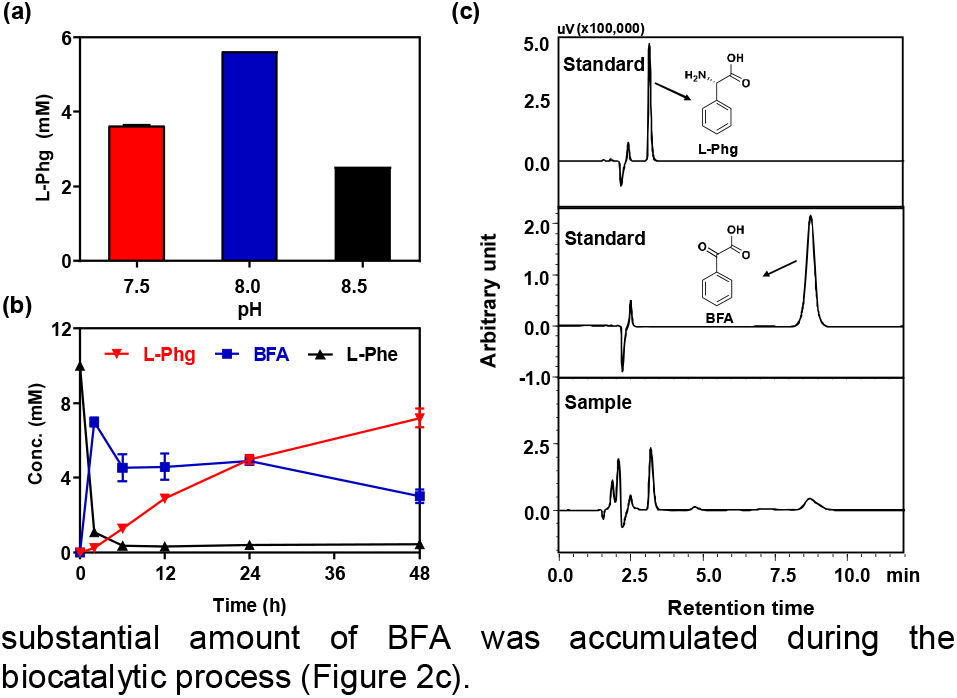
Characterization of AT-dependent route for L-Phg synthesis. (a) The efficiency of biotransforming L-Phe (10 mM) to L-Phg under different pH conditions. (b) Time course of biotransforming L-Phe (10 mM) to L-Phg at pH 8.0. (c) Representative HPLC result showing L-Phg produced by the recombinant *E. coli* strain Ec-Phg1.0. All experiments mentioned above were carried out in 200 mM KP buffer with 10 g cdw L^−1^ recombinant *E. coli* at 30°C. Data represent the mean value with standard deviations from triplicate of experiments.

According to the literature, LAAD from *P. mirabilis* showed a broad substrate activity on L-amino acids, such as L-His, L-Arg, L-Phe and so on^[19]^. We hypothesized that LAAD might deaminate L-Phg, which leads to a futile catalytic cycle and results in the accumulation of BFA. To corroborate this hypothesis, we further carried out the L-Phg stability test. When 10 mM L-Phg was treated with the recombinant *E. coli* expressing LAAD, no appreciable degradation of L-Phg was observed even after 48 h (Supplementary Figure S1). Therefore, we suspected that the transamination step catalyzed by AT/GluDH might be the bottleneck limiting the L-Phg production. As it was reported that the production of 34 mM L-Phg from 40 mM L-Phe was achieved in *E. coli* containing eight reaction steps with AT/GluDH and there was no accumulation of BFA in the experimental process^[12]^, the AT activity is unlikely to be rate-limiting. However, LAAD from *P. mirabilis* also effectively deaminates L-glutamate to α-ketoglutarate ^[20]^. Thus, we inferred that the amine donor of L-glutamate was degraded by LAAD, thereby leading to the accumulation of BFA and poor yield of L-Phg.

To overcome the low catalytic efficiency of AT/GluDH-mediated route toward L-Phg production, we next sought to find an alternative strategy to skip the use of amine donor of L-glutamate. From the literature, the preparation of L-Phg in *E. coli* could be realized by expressing an amino acid dehydrogenase from *Bacillus clausii* (BcAADH) with optimal pH around 9.5-10.5 ^[21]^, LeuDH from *Exiguobacterium sibiricum* with optimal pH around 10.0, ^[22]^ and LeuDH from *B. cereus* with optimal pH around 8.0 ^[16]^. In this study, we decided to replace AT/GluDH with LeuDH from *B. cereus*, which is more compatible with our establishment reaction condition (Scheme 1). As shown in Figure 3a, we further constructed the recombinant *E. coli* strain

**Figure 3.**
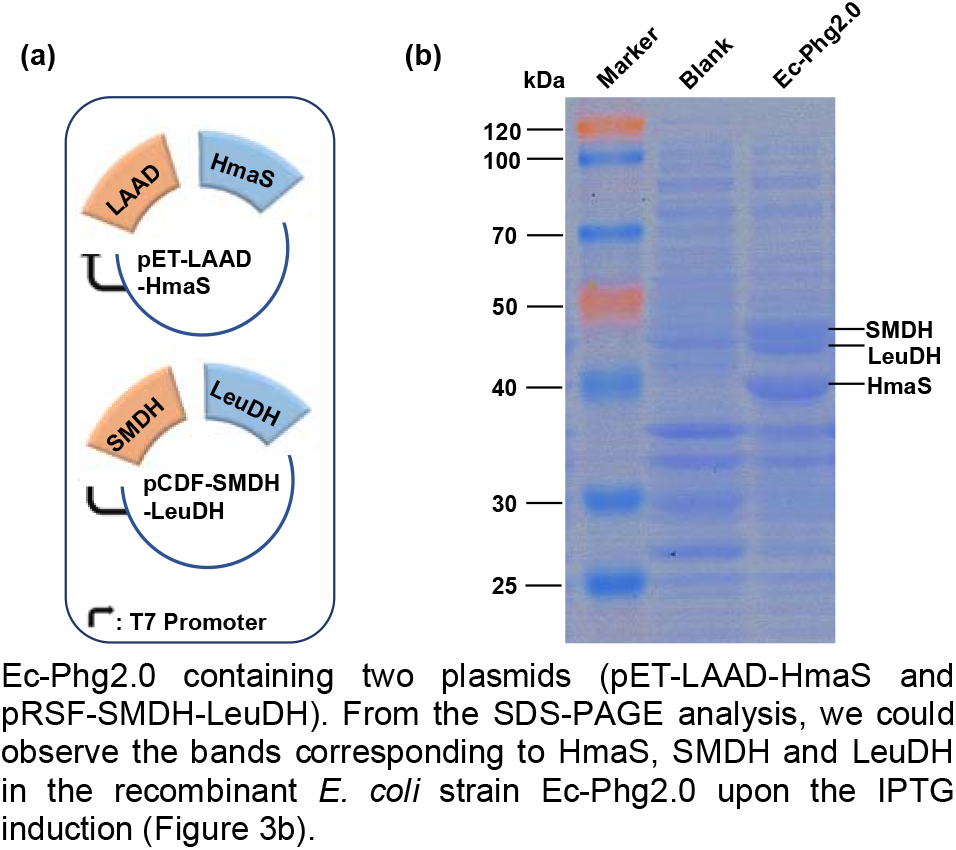
Plasmid design of leucine dehydrogenase (LeuDH)-dependent route and SDS-PAGE analysis of protein expression. (a) The recombinant *E. coli* strain Ec-Phg2.0 containing two plasmids (pET-LAAD-HmaS and pRSF-SMDH-LeuDH). (b) SDS-PAGE analysis of the recombinant *E. coli* strain Ec-Phg2.0. Blank: *E. coli* BL21 (DE3) harboring empty plasmids.

Ec-Phg2.0 containing two plasmids (pET-LAAD-HmaS and pRSF-SMDH-LeuDH). From the SDS-PAGE analysis, we could observe the bands corresponding to HmaS, SMDH and LeuDH in the recombinant *E. coli* strain Ec-Phg2.0 upon the IPTG induction (Figure 3b).

As can be seen from Figure 4a and Table S2, when 10 mM L-Phe was used as the substrate, 9.92 ± 0.39 mM L-Phg was produced after 12 h. Interestingly, the problem of BFA accumulation was addressed by replacing AT/GluDH module with LeuDH from *B. cereus* (Figure 4c). To compare the catalytic efficiency of our system with that of the recombinant *E. coli* expressing 12 genes ^[12]^, we also examined high substrate concentration of 40 mM L-Phe. When 40 mM L-Phe was fed to the recombinant *E. coli*, approximately 39.97 ± 3.84 mM (6.04 ± 0.58 g/L) L-Phg was obtained after 12 h (Figure 4b, Table S3), which corresponds to >99.9% conversion.

**Figure 4.**
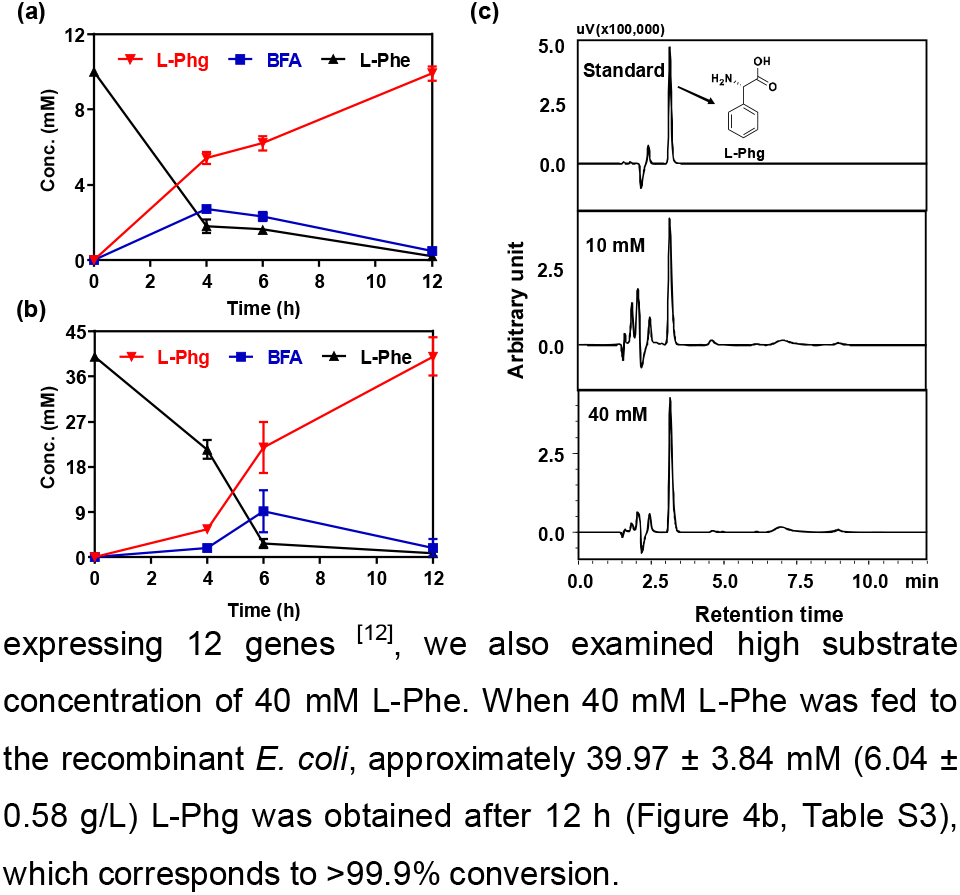
Characterization of LeuDH-dependent route for L-Phg synthesis. Time course of biotransforming L-Phe (10 mM) to L-Phg. (b) Time course of bioconverting L-Phe (40 mM) to L-Phg. (c) Representative HPLC result showing L-Phg produced by the recombinant *E. coli* strain Ec-Phg2.0. All experiments mentioned above were carried out in 200 mM KP buffer with 10 g cdw L^−1^ recombinant *E. coli* at 30°C. Data represent the mean value with standard deviations from triplicate of experiments.

In summary, we have developed two artificial enzymatic cascades to synthesize enantiopure amino acid of L-Phg from biobased L-Phe. The first enzymatic cascade comprises co-expressing five enzymes in the recombinant *E. coli* strain Ec-Phg1.0 harboring three corresponding plasmids (pET-LAAD-HmaS, pRSF-SMDH-AT and pACYC-GluDH). However, under the optimal pH condition, only 7.21 ± 0.15 mM L-Phg was obtained from 10 mM L-Phe after 48 h and the accumulation of intermediate product BFA was observed. Further experiments revealed that LAAD from *P. mirabilis* could not deaminate L-Phg (Supplementary Figure S1), indicating that the transamination step is rate-limiting. Next, we attempted to use LeuDH from *B. cereus* to replace AT/GluDH and the recombinant *E. coli* strain Ec-Phg2.0 harboring two corresponding plasmids (pET-LAAD-HmaS and pRSF-SMDH-LeuDH) was constructed. The four-step enzymatic cascade could efficiently convert 40 mM of L-Phe to 39.97 ± 3.84 mM (6.04 ± 0.58 g/L) L-Phg, reaching >99.9% conversion after 12 h. Although we did not further optimize and scale-up the biocatalytic process, it would be possible to achieve even higher L-Phg titer as more active enzyme alternatives are available for L-Phg synthesis from mandelate ^[21–22, 25]^. Based on these findings, the four-step enzymatic cascade outperformed all the previously established biocatalytic systems for L-Phg synthesis from biobased L-Phe ^[12], [24]^. Since there was no appreciable accumulation of intermediates for *E. coli* strain Ec-Phg2.0, our biocatalytic method would greatly simplify the product separation process, with a great potential for future industrial applications.

## Acknowledgements

This work was supported by Xiamen University under grant no. 0660-X2123310 and ZhenSheng Biotech, China.

## Notes

### Competing Interest Statement

The authors have declared no competing interest.

