## Supplemental materials for "A Four-Step Enzymatic Cascade for Efficient Production of L-Phenylglycine from Biobased L-Phenylalanine"

### 1 Experimental Procedures

#### 1.1 Chemicals, biochemicals and medium

**All chemicals were purchased from commercial suppliers and used without further purification:**
Chemicals purchased from Sigma-Aldrich (St. Louis, MO, USA): D-glucose (99%), L-glutamate (98.5%), L-phenylalanine (≥98%), mandelic acid (99%), L-phenylglycine (99%)
Chemicals purchased from Inalco (San Luis Obispo, CA, USA): ampicillin sodium salt (>84.5%), kanamycin sulfate salt (>75%), chloramphenicol (>85%) Isopropyl β-D-1-thiogalactopyranoside (IPTG, >99%).
Chemicals purchased from Sinopharm Chemical Reagent (Shanghai, China): glycerol (>99%), HCl (37%), NaCl (≥99.5%), $K_2HPO_4•3H_2O$ (≥99%), $KH_2PO_4$(≥99.5%), $NH_4Cl$ (≥99.5%), $NH_3•H_2O$ (25~28%).
Chemicals purchased from Shanghai Aladdin Biochemical Technology (Shanghai, China): trifluoroacetic acid (HPLC, ≥99.5%).
Chemicals purchased from Shanghai Macklin Biochemical Co., Ltd (Shanghai, China): benzoylformic acid (>95.0%)
Chemicals purchased from ANPEL laboratory Technologies Inc. (Shanghai, China): acetonitrile (HPLC, ≥99.9%).
**Biochemicals and kits were purchased from commercial suppliers:**
GenScript (Nanjing, Jiangsu, China): DNA oligonucleotides and genes.
New England Biolabs (Beverly, MA, USA): Phusion DNA polymerase, dNTP mixture, restriction enzymes, T4 DNA ligase.
Biomer Technology (Runcorn, Cheshire, UK): plasmid miniprep kit, gel extraction kit, DNA purification kit.
Sangon Biotech (Shanghai, China): LB medium, tryptone, yeast extract, agar.
TransGene (Beijing, China): *Blue Plus*® II Protein Marker (14-120 kDa)
Luria-Bertani medium was used for molecular cloning and the initial seed culture.
Terrific broth (TB) containing $K_2HPO_4$ (12.54 g/L), $KH_2PO_4$ (2.32 g/L), tryptone (12 g/L), glycerol (0.4%) and yeast extract (24 g/L) was used for culturing the *Escherichia coli* cells (50-200 mL) for biotransformation.

#### 1.2 Construction of plasmids and strains

All the genes were PCR amplified using High-fidelity Phusion Polymerase from New England Biolabs (Beverly, MA, USA). The genes encoding LAAD from *P. Mirabilis* and HmaS from *A. orientalis* were synthesized and codon optimized for *E. coli* by GenScript (Nanjing, Jiangsu, China). The genes encoding SMDH and LeuDH were obtained from *P. putida* and *B. cereus,* respectively. The genes encoding AT and GluDH were obtained from *E. coli*. All the primers used in the present study are shown in Supplementary Table S4.
All plasmids pET-LAAD-HmaS, pRSF-SMDH-AT, pACYC-GluDH and pRSF-SMDH-LeuDH were constructed via the standard T4 ligase mediated approach. After verification by restriction enzyme digestion and DNA sequencing from Sangon Biotech (Shanghai, China), the plasmids were transformed into *E. coli* BL21 (DE3) via standard electroporation or heat-shock approach. All enzymes (*Bsa*I-HF, *Bam*HI, *Xho*I, T4 DNA ligase) were purchased from New England Biolabs (Beverly, MA, USA). All the plasmids and strains used in the present study are listed in Supplementary Table S5.

#### 1.3 Recombinant *E. coli* strain cultivation

The recombinant *E. coli* strain Ec-Phg1.0 harboring three corresponding plasmids (pET-LAAD-HmaS, pRSF-SMDH-AT and pACYC-GluDH) was inoculated in 2 mL Luria-Bertani medium (LB, tryptone 10 g $L^{-1}$, NaCl 10 g $L^{-1}$, yeast extract 5 g $L^{-1}$) with appropriate antibiotics (33 μg $mL^{-1}$ ampicillin, 17 μg $mL^{-1}$ kanamycin and 11 μg $mL^{-1}$ chloramphenicol). The *E. coli* cells were cultivated at 37°C and 250 rpm for overnight. Then, 1 mL fresh overnight culture was inoculated into a 250 mL flask containing 75 mL Terrific Broth ($KH_2PO_4$ 2.32 g $L^{-1}$, $K_2HPO_4•3H_2O$ 12.54 g $L^{-1}$, tryptone 12 g $L^{-1}$, yeast extract 24 g $L^{-1}$ and glycerol 0.4%) with appropriate antibiotics and the cells were allowed to grow at 37°C and 250 rpm until the optical density at 600 nm (OD600) reached 0.8~0.9. IPTG (isopropyl *β*-D-1-thiogalactopyranoside, Inalco, San Luis Obispo, CA, USA) was added at a final concentration of 1 mM to induce protein expression. After further cultivation at 20°C for another 18 h, the cells were collected by centrifugation for 5 min at 7000 rpm and 4°C.

The cell pellets were washed once with ice-cold $ddH_2O$, and resuspended to a final cell density of 40 g cdw $L^{-1}$ with 200 mM potassium phosphate buffer (KP buffer, $K_2HPO_4$•$3H_2O$ 42.90 g $L^{-1}$, $KH_2PO_4$ 1.63 g $L^{-1}$, pH 8.0). The recombinant *E. coli* strain Ec-Phg2.0 harboring two corresponding plasmids (pET-LAAD-HmaS and pRSF-SMDH-LeuDH) was cultivated in a similar manner as described above.

#### 1.4 Biotransformation procedure

For L-Phg production with the recombinant strain Ec-Phg1.0, a 2 mL reaction mixture was prepared: 10 mM L-Phe, 50 mM L-glutamate, 200 mM $NH_4Cl/NH_3$, 2% glucose, Ec-Phg1.0 cell suspension (10 g cdw $L^{-1}$) in KP buffer (200 mM, pH 8.0). For L-Phg production with the recombinant strain Ec-Phg2.0, a 2 mL reaction mixture was prepared: 10 mM or 40 mM L-Phe, 200 mM $NH_4Cl/NH_3$, 2% glucose, Ec-Phg2.0 cell suspension (10 g cdw $L^{-1}$) in KP buffer (200 mM, pH 8.0). For investigating the LAAD activity on L-Phg, the recombinant *E. coli* strain harboring pET-LAAD-HmaS was fed with 10 mM L-Phg. 1 mL reaction mixture contains: 10 mM L-Phg, cell suspension 10 g cdw $L^{-1}$ in KP buffer (200 mM, pH 8.0). The condition for biotransformation was set at 30°C and 250 rpm on a rotary shaking incubator. Samples were periodically collected and analyzed by high performance liquid chromatography (HPLC) (Shimadzu prominence LC-20A, Japan).

#### 1.5 Analytical methods

The reaction mixtures were first diluted by the solution containing 0.1% trifluoroacetate (Aladdin, Shanghai, China) and 10% acetonitrile (ANPEL, Shanghai, China), and then centrifuged to remove the cell pellet. The supernatant was filtered before used for HPLC analysis by the Shimadzu prominence LC-20A system equipped with a reversed-phase column Shimadzu C18 column (150 mm × 4.6 mm × 5 μm) and a photodiode array detector (DAD). The mobile phase containing 90% water with 0.1% trifluoroacetate and 10% acetonitrile was used to analyse L-Phg, L-Phe and BFA. The detection wave length of L-Phg is 210 nm and the detection wave length of L-Phe and BFA is 254 nm. The flow rate was set at 1 mL $min^{-1}$, and the column temperature was maintained at 40°C. The retention times for determining the analytes are as the following: L-Phg is 3.2 min; L-Phe, 6.2 min; BFA, 8.7 min. The standard samples of L-Phg, L-Phe and BFA were diluted to 1 mM, 0.5 mM, 0.25 mM, and 0.125 mM in equal gradient to prepare the calibration curves.

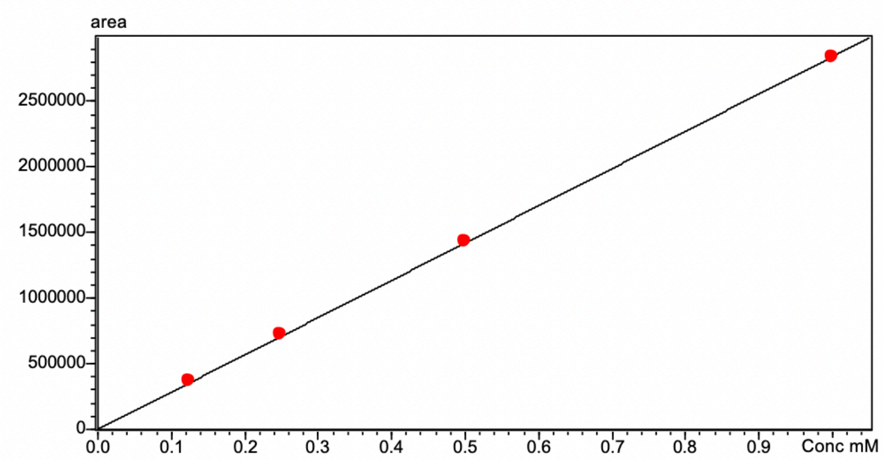


**The calibration curve of L-Phg.** The calibration curve formula of L-Phg is Y=2.83953e+006X+0, $R^2$=0.9999.

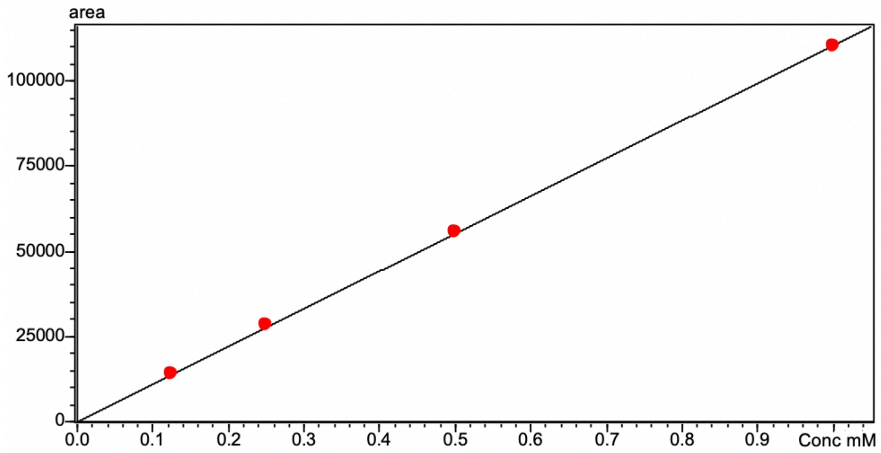


**The calibration curve of L-Phe.** The calibration curve formula of L-Phe is Y=110359X+0, $R^2$=0.9999.

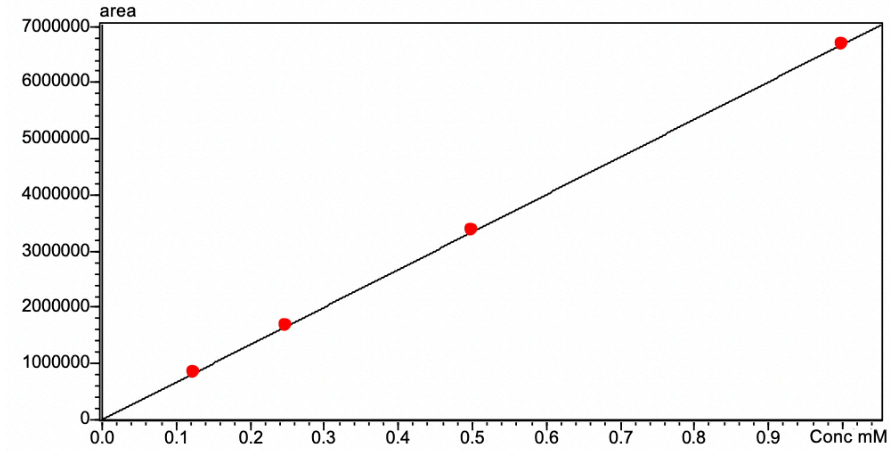


**The calibration curve of BFA.** The calibration curve formula of BFA is Y=6.68108e+006X+0, $R^2$=0.9999.

#### 1.6 Gene sequences

*LAAD* gene from *P. mirabilis* was codon optimized and synthesized by GenScript. The optimized DNA sequence is:

>LAAD

ATGAACATTTCACGTCGCAAGCTGCTGCTGGGTGTTGGTGCTGCTGGTGTGTTGGCGGGTGGTGCAGCTCTGGTTCCAATGGTG
CGTCGTGATGGTAAATTTGTTGAAGCAAAGAGCCGTGCGAGCTTCGTGGAAGGTACCCAAGGTGCGCTGCCGAAAGAAGCTGA
CGTTGTGATTATCGGTGCTGGTATTCAGGGTATCATGACCGCTATTAATCTGGCAGAACGTGGTATGAGCGTTACCATTCTGGAA
AAGGGTCAAATCGCAGGTGAACAGAGCGGTCGTGCGTACAGCCAAATTATCAGCTATCAGACCAGCCCGGAAATTTTTCCGCTG
CATCACTACGGTAAAATCCTGTGGCGTGGTATGAACGAAAAGATTGGTGCGGATACCAGCTATCGTACCCAAGGTCGTGTTGAA
GCTCTGGCAGATGAAAAAGCACTGGACAAGGCGCAGGCTTGGATCAAAACCGCGAAGGAAGCTGCAGGTTTTGACACCCCGCT
GAATACCCGTATTATCAAAGGTGAAGAACTGAGCAACCGTCTGGTTGGTGCTCAAACCCCGTGGACCGTGGCTGCTTTCGAAGA
AGATAGCGGTAGCGTTGACCCGGAAACCGGTACCCCGGCACTGGCTCGTTACGCTAAACAGATTGGTGTTAAGATCTATACCAA
CTGCGCTGTGCGTGGTATTGAAACCGCGGGTGGTAAAATCAGCGATGTTGTGAGCGAAAAAGGTGCGATCAAGACCAGCCAAG
TGGTGCTGGCGGGTGGTATTTGGAGCCGTCTGTTTATGGGTAATATGGGTATTGACATCCCGACCCTGAACGTTTACCTGAGCC
AACAACGTGTTAGCGGTGTGCCAGGTGCGCCGCGTGGTAATGTGCATCTGCCGAACGGTATCCACTTTCGTGAACAAGCTGATG
GTACCTATGCTGTTGCACCGCGTATTTTCACCAGCAGCATCGTGAAAGACAGCTTTCTGCTGGGTCCGAAGTTCATGCATCTGCT
GGGTGGTGGTGAACTGCCGCTGGAATTTTCTATCGGTGAAGACCTGTTTAATAGCTTCAAAATGCCGACCAGCTGGAACCTGGA
CGAAAAGACCCCGTTTGAACAATTCCGTGTTGCGACCGCTACCCAAAATACCCAGCACCTGGATGCAGTTTTTCAGCGTATGAAA
ACCGAATTTCCGGTGTTCGAAAAGAGCGAAGTTGTGGAACGTTGGGGTGCTGTTGTGAGCCCGACCTTCGACGAACTGCCGATT
ATCAGCGAAGTTAAGGAATACCCGGGTCTGGTTATTAACACCGCTACCGTGTGGGGTATGACCGAAGGTCCGGCAGCGGGTGA
AGTTACCGCAGATATTGTGATGGGTAAAAAGCCGGTTATTGATCCGACCCCGTTTAGTTTGGATCGTTTTAAGAAGTAA

*HmaS* gene from *A. orientalis* was codon optimized and synthesized by GenScript. The optimized DNA sequence is:

> HmaS

ATGCAGAATTTCGAAATCGACTATGTTGAGATGTATGTGGAGAATCTGGAAGTGGCGGCTTTCTCATGGGTCGACAAGTACGCAT
TCGCCGTGGCTGGTACGAGCCGTTCAGCGGACCACAGGAGCATCGCGTTGAGACAAGGGCAGGTGACACTTGTTCTTACTGAA
CCGACGTCAGATAGACATCCGGCCGCCGCGTATTTGCAGACTCATGGCGATGGTGTAGCGGATATTGCTATGGCCACCTCCGA
CGTTGCTGCGGCCTACGAGGCGGCAGTACGTGCCGGTGCTGAAGCTGTTAGAGCACCAGGGCAACACAGTGAAGCTGCTGTGA
CGACCGCGACCATAGGTGGGTTTGGAGATGTCGTCCATACTCTGATCCAAAGGGACGGCACTAGCGCTGAATTACCCCCCGGA
TTTACCGGCTCCATGGACGTTACGAACCATGGTAAAGGAGACGTAGATCTTCTGGGGATAGATCACTTTGCGATATGTCTTAACG
CCGGAGATCTGGGACCTACTGTGGAATACTACGAGAGAGCTTTAGGTTTTAGGCAAATATTTGACGAGCATATAGTTGTTGGCGC
TCAGGCGATGAACTCAACTGTCGTGCAAAGTGCGAGTGGGGCGGTAACACTAACCTTGATAGAACCCGATCGTAATGCCGACCC
CGGGCAAATTGATGAGTTCCTAAAAGATCACCAGGGTGCGGGTGTGCAGCACATCGCCTTTAATTCTAATGACGCTGTGCGTGC
AGTGAAAGCACTGTCAGAGAGAGGGGTCGAGTTTTTGAAAACCCCGGGGGCGTATTACGATCTTCTAGGAGAGAGAATAACCCT
GCAAACGCATAGTCTGGATGATTTGAGAGCGACCAATGTTTTGGCAGATGAGGATCACGGAGGTCAACTTTTTCAAATCTTCACA
GCGAGTACGCACCCAAGACACACCATTTTTTTTGAAGTCATCGAAAGACAAGGAGCGGGCACTTTTGGTTCCAGTAATATCAAGG
CTTTATATGAGGCAGTGGAACTAGAACGTACAGGTCAAAGTGAGTTTGGAGCCGCGAGGCGTTAG

*mdlB* gene was amplified from *P. putida*

>mdlB:

ATGAGCCGGAATCTCTTTAACGTTGAGGACTATCGCAAGCTCGCGCAAAAGCGCTTGCCAAAGATGGTGTACGACTATCTGGAA
GGTGGAGCTGAAGACGAATACGGGGTGAAACACAACCGCGACGTCTTCCAGCAATGGCGATTCAAACCGAAGCGGCTGGTAGA
TGTCAGCCGCCGCAGCCTTCAAGCGGAAGTACTTGGAAAGAGGCAGTCAATGCCTCTCTTGATTGGCCCAACTGGGCTGAATGG
TGCGCTCTGGCCTAAAGGGGATCTCGCCCTTGCTCAAGCGGCAACCAAGGCCGGCATCCCGTTCGTGCTGTCGACCGCCTCCA
ACATGTCCATTGAAGACCTCGCTCGTCAGTGTGATGGCGATCTATGGTTCCAGCTCTATGTGATCCACCGAGAGATCGCGCAGG
GGATGGTGCTCAAAGCCCTGCACTCCGGTTACACCACGCTGGTACTCACAACAGACGTCGCGGTTAATGGCTATCGCGAGCGA
GACCTGCACAACCGATTCAAGATGCCGATGAGCTACACCCCAAAGGTGATGCTGGACGGATGCCTGCATCCACGCTGGTCGCT
CGATCTGGTGCGCCACGGCATGCCGCAACTGGCCAACTTCGTCAGCAGTCAAACGTCCAGCTTAGAGATGCAGGCAGCATTGA
TGAGCCGCCAAATGGATGCCAGTTTCAACTGGGAGGCATTGAGATGGCTGCGTGACCTCTGGCCGCACAAACTCCTCGTAAAG
GGGTTGCTCAGTGCTGAGGACGCCGATCGGTGCATCGCTGAAGGTGCCGACGGCGTGATCCTGTCAAACCACGGCGGTCGCC
AACTCGATTGCGCGGTATCGCCAATGGAAGTTTTGGCTCAATCGGTAGCGAAAACTGGAAAACCAGTGCTTATCGATAGCGGCT
TCCGACGGGGTTCGGACATCGTTAAAGCGCTTGCGCTAGGTGCTGAGGCTGTACTCCTGGGACGTGCAACTTTGTATGGCCTTG
CAGCACGAGGTGAAACGGGTGTTGGCGAGGTGCTAACCCTCCTCAAAGCGGATATCGACCGCACTCTGGCCCAGATTGGATGC
CCTGACATCACCTCCCTTTCTCCTGATTACCTCCAAAGCGAGGGAGTGACTAACACCGCTCCAGTCGATCACCTCATTGGTAAAG
GAACACACGCATGA

*ilvE* gene from *E. coli*

> *ilvE*

ATGACCACGAAGAAAGCTGATTACATTTGGTTCAATGGGGAGATGGTTCGCTGGGAAGACGCGAAGGTGCATGTGATGTCGCAC
GCGCTGCACTATGGCACTTCGGTTTTTGAAGGCATCCGTTGCTACGACTCGCACAAAGGACCGGTTGTATTCCGCCATCGTGAG
CATATGCAGCGTCTGCATGACTCCGCCAAAATCTATCGCTTCCCGGTTTCGCAGAGCATTGATGAGCTGATGGAAGCTTGTCGT
GACGTGATCCGCAAAAACAATCTCACCAGCGCCTATATCCGTCCGCTGATCTTCGTCGGTGATGTTGGCATGGGAGTAAACCCG
CCAGCGGGATACTCAACCGACGTGATTATCGCTGCTTTCCCGTGGGGAGCGTATCTGGGCGCAGAAGCGCTGGAGCAGGGGAT
CGATGCGATGGTTTCCTCCTGGAACCGCGCAGCACCAAACACCATCCCGACGGCGGCAAAAGCCGGTGGTAACTACCTCTCTT
CCCTGCTGGTGGGTAGCGAAGCGCGCCGCCACGGTTATCAGGAAGGTATCGCGCTGGATGTGAACGGTTATATCTCTGAAGGC
GCAGGCGAAAACCTGTTTGAAGTGAAAGATGGTGTGCTGTTCACCCCACCGTTCACCTCCTCCGCGCTGCCGGGTATTACCCGT
GATGCCATCATCAAACTGGCGAAAGAGCTGGGAATTGAAGTACGTGAGCAGGTGCTGTCGCGCGAATCCCTGTACCTGGCGGA
TGAAGTGTTTATGTCCGGTACGGCGGCAGAAATCACGCCAGTGCGCAGCGTAGACGGTATTCAGGTTGGCGAAGGCCGTTGTG
GCCCGGTTACCAAACGCATTCAGCAAGCCTTCTTCGGCCTCTTCACTGGCGAAACCGAAGATAAATGGGGCTGGTTAGATCAAG
TTAATCAATAA

*GluDH* gene from *E. coli*

>GluDH

ATGGATCAGACATATTCTCTGGAGTCATTCCTCAACCATGTCCAAAAGCGCGACCCGAATCAAACCGAGTTCGCGCAAGCCGTTC
GTGAAGTAATGACCACACTCTGGCCTTTTCTTGAACAAAATCCAAAATATCGCCAGATGTCATTACTGGAGCGTCTGGTTGAACC
GGAGCGCGTGATCCAGTTTCGCGTGGTATGGGTTGATGATCGCAACCAGATACAGGTCAACCGTGCATGGCGTGTGCAGTTCA

GCTCTGCCATCGGCCCGTACAAAGGCGGTATGCGCTTCCATCCGTCAGTTAACCTTTCCATTCTCAAATTCCTCGGCTTTGAACA
AACCTTCAAAAATGCCCTGACTACTCTGCCGATGGGCGGTGGTAAAGGCGGCAGCGATTTCGATCCGAAAGGAAAAAGCGAAG
GTGAAGTGATGCGTTTTTGCCAGGCGCTGATGACTGAACTGTATCGCCACCTGGGCGCGGATACCGACGTTCCGGCAGGTGAT
ATCGGGGTTGGTGGTCGTGAAGTCGGCTTTATGGCGGGGATGATGAAAAAGCTCTCCAACAATACCGCCTGCGTCTTCACCGGT
AAGGGCCTTTCATTTGGCGGCAGTCTTATTCGCCCGGAAGCTACCGGCTACGGTCTGGTTTATTTCACAGAAGCAATGCTAAAAC
GCCACGGTATGGGTTTTGAAGGGATGCGCGTTTCCGTTTCTGGCTCCGGCAACGTCGCCCAGTACGCTATCGAAAAAGCGATG
GAATTTGGTGCTCGTGTGATCACTGCGTCAGACTCCAGCGGCACTGTAGTTGATGAAAGCGGATTCACGAAAGAGAAACTGGCA
CGTCTTATCGAAATCAAAGCCAGCCGCGATGGTCGAGTGGCAGATTACGCCAAAGAATTTGGTCTGGTCTATCTCGAAGGCCAA
CAGCCGTGGTCTCTACCGGTTGATATCGCCCTGCCTTGCGCCACCCAGAATGAACTGGATGTTGACGCCGCGCATCAGCTTATC
GCTAATGGCGTTAAAGCCGTCGCCGAAGGGGCAAATATGCCGACCACCATCGAAGCGACTGAACTGTTCCAGCAGGCAGGCGT
ACTATTTGCACCGGGTAAAGCGGCTAATGCTGGTGGCGTCGCTACATCGGGCCTGGAAATGGCACAAAACGCTGCGCGCCTGG
GCTGGAAAGCCGAGAAAGTTGACGCACGTTTGCATCACATCATGCTGGATATCCACCATGCCTGTGTTGAGCATGGTGGTGAAG
GTGAGCAAACCAACTACGTGCAGGGCGCGAACATTGCCGGTTTTGTGAAGGTTGCCGATGCGATGCTGGCGCAGGGTGTGATT
TAA

*LeuDH* gene from *B. cereus*

> LeuDH

ATGACATTAGAAATCTTCGAATACTTAGAAAAATATGATTATGAGCAAGTAGTATTTTGTCAAGATAAAGAATCTGGTTTAAAAGCA
ATTATTGCAATTCATGATACAACACTTGGACCGGCTCTTGGTGGAACAAGAATGTGGACATATGATTCTGAAGAAGCGGCGATTG
AAGATGCATTGCGTCTTGCAAAAGGGATGACATACAAAAACGCAGCAGCTGGTTTAAACTTAGGTGGTGCGAAAACAGTAATTAT
CGGTGATCCTCGTAAAGATAAGAGCGAAGCAATGTTCCGTGCACTAGGACGTTATATCCAAGGACTAAACGGACGTTACATTACA
GCTGAAGATGTTGGTACAACAGTAGATGATATGGATATTATCCATGAAGAAACTGACTTTGTAACAGGTATCTCACCATCATTCGG
TTCTTCTGGTAACCCATCTCCGGTAACTGCATACGGTGTTTACCGTGGTATGAAAGCAGCTGCAAAAGAAGCTTTCGGTACTGAC
AATTTAGAAGGAAAAGTAATTGCTGTTCAAGGCGTTGGTAACGTAGCATATCACCTATGCAAACATTTACACGCTGAAGGAGCAA
AATTAATTGTTACAGATATTAATAAAGAAGCTGTACAACGTGCTGTAGAAGAATTCGGTGCATCAGCAGTTGAACCAAATGAAATT
TACGGTGTTGAATGCGATATTTACGCACCATGTGCACTAGGCGCAACAGTTAATGATGAAACTATTCCACAACTTAAAGCAAAAGT
AATCGCAGGTTCTGCGAATAACCAATTAAAAGAAGATCGTCATGGTGACATCATTCATGAAATGGGTATTGTATACGCACCAGATT
ATGTAATTAATGCAGGTGGCGTAATTAACGTAGCAGACGAATTATATGGATACAATAGAGAACGTGCACTAAAACGTGTTGAGTCT
ATTTATGACACGATTGCAAAAGTAATCGAAATTTCAAAACGCGATGGCATAGCAACTTATGTAGCGGCAGATCGTCTAGCTGAAG
AGCGCATTGCAAGCTTGAAGAATTCTCGTAGCACTTACTTACGCAACGGTCACGATATTATTAGCCGTCGCTAA

#### 2 Supporting Figure

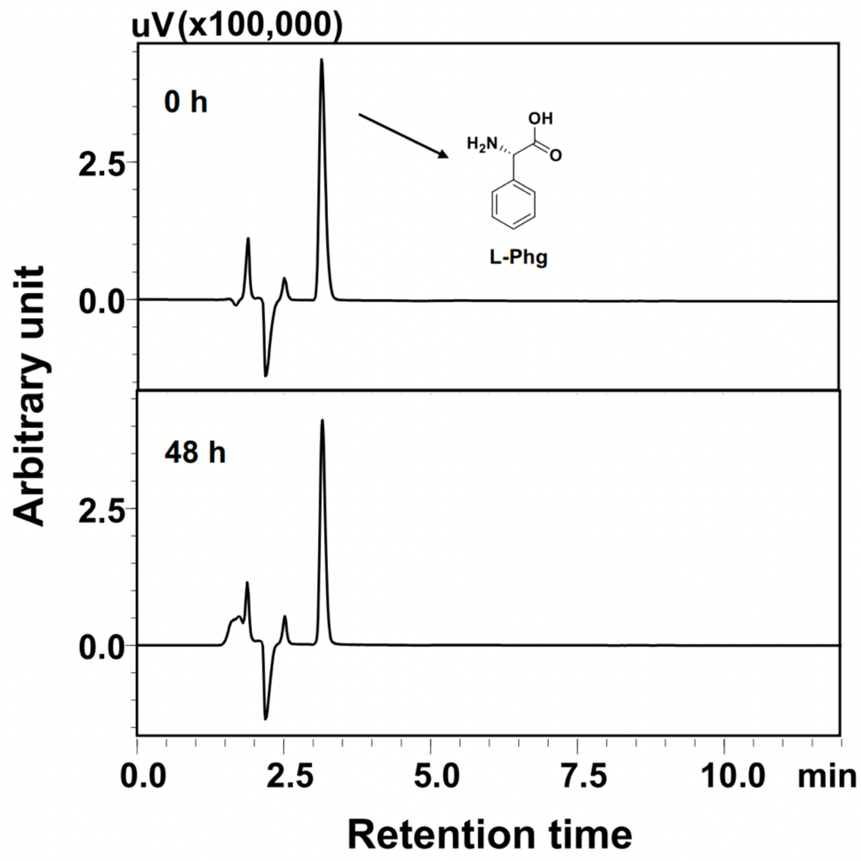


**Figure S1. L-Amino acid deaminase (LAAD) from *P. mirabilis* could not deaminate L-phenylglycine.** Samples from the recombinant *E. coli* strain of Ec-LAAD (0 h and 48 h) were subjected to HPLC analysis. L-Phenylglycine was stably maintained after 48 h treatment with the recombinant *E. coli* strain expressing LAAD from *P. mirabilis*.

#### 3 Supporting Tables

**Table S1. Biotransforming L-Phe (10 mM) to L-Phg via the AT-dependent route.**

| Time (h) | L-Phg (mM) | BFA (mM) | L-Phe (mM) |
|---|---|---|---|
| 2 | 0.22±0.01 | 6.99±0.25 | 1.09±0.05 |
| 6 | 1.28±0.10 | 4.53±0.74 | 0.35±0.02 |
| 12 | 2.89±0.17 | 4.59±0.70 | 0.34±0.09 |
| 24 | 4.97±0.25 | 4.89±0.24 | 0.42±0.08 |
| 48 | 7.21±0.51 | 3.01±0.36 | 0.44±0.11 |

**Table S2. Biotransforming L-Phe (10 mM) to L-Phg via the LeuDH-dependent route.**

| Time (h) | L-Phg (mM) | BFA (mM) | L-Phe (mM) |
|---|---|---|---|
| 4 | 5.74±0.32 | 2.71±0.15 | 1.80±0.36 |
| 6 | 6.21±0.39 | 2.31±0.26 | 1.61±0.17 |
| 12 | 9.92±0.39 | 0.47±0.40 | 0.21±0.07 |

**Table S3. Biotransforming L-Phe (40 mM) to L-Phg via the LeuDH-dependent route.**

| Time (h) | L-Phg (mM) | BFA (mM) | L-Phe (mM) |
|---|---|---|---|
| 4 | 5.60±0.18 | 1.87±0.31 | 21.43±1.88 |
| 6 | 21.83±5.02 | 9.11±4.15 | 2.73±0.97 |
| 12 | 39.97±3.84 | 1.84±1.78 | 0.68±0.59 |

**Table S4. List of primers used in this study.**

| Name of primers [a] | Sequence |
|---|---|
| LAAD_fwd | TTGGTCTCGGATCCGATGAACATTTCACGTCGCAAGC |
| LAAD_rev | TTGGTCTCATCCTTTACTTCTTAAAACGATCCAAAC |
| HmaS_fwd | TTGGTCTCAAGGAGATATATTATGCAGAATTTCGAAATC |
| HmaS_rev | TTGGTCTCCTCGAGCTAACGCCTCGCGGCTCC |
| mdlB_fwd | TTGGTCTCGGATCCGATGAGCCGGAATCTCTTTAAC |
| mdlB_OE_rev | GTTGTGCAGGTCcCGCTCGCG |
| mdlB_OE_fwd | CGCGAGCGGGACCTGCACAAC |
| mdlB_rev | TTGGTCTCATCCTTCATGCGTGTGTTCCTTTAC |
| ilvE_fwd | TTGGTCTCAAGGAGATATATAATGACCACGAAGAAAGCTG |
| ilvE_rev | TTGGTCTCCTCGAGTTATTGATTAACTTGATCTAAC |
| GluDH_fwd | CGCGGATCCGATGGATCAGACATATTCTC |
| GluDH_rev | AGAGACTCGAGTTAAATCACACCCTGCGCC |
| LeuDH_fwd | TTGGTCTCAAGGAGATAACTTTATGACATTAGAAATCTTCG |
| LeuDH_rev | TTGGTCTCCTCGAGTTAGCGACGGCTAATAATATCG |

[a] The oligonucleotides were synthesized by GenScript.

**Table S5. List of plasmids and strains used in this study.**

| Name | Features | References |
|---|---|---|
| **Plasmids** | | |
| pETDuet-1 | $Ap^R$, lacI, T7lac | Novagen |
| pRSFDuet-1 | $Km^R$, lacI, T7lac | Novagen |
| pACYCDuet-1 | $Cm^R$, lacI, T7lac | Novagen |
| pET-LAAD-HmaS | pETDuet-1 harboring LAAD (L-amino acid deaminase from *P. mirabilis*) and HmaS (hydroxymandelate synthase from *A. orientalis*) | This study |
| pRSF-SMDH-AT | pRSFDuet-1 harboring SMDH ((*S*)-mandelate dehydrogenase from *P. putida*) and AT (aminotransferase from *E. coli*) | This study |
| pACYC-GluDH | pACYCDuet-1 harboring GluDH (glutamate dehydrogenase from *E. coli)* | This study |
| pRSF-SMDH-LeuDH | pRSFDuet-1 harboring SMDH ((*S*)-mandelate dehydrogenase from *P. putida*) and LeuDH (leucine dehydrogenase from *B. cereus*) | This study |
| **Strains** | | |
| *E. coli* TOP 10 | For cloning purpose | Novagen |
| *E. coli* BL21 (DE3) | For protein overexpression | Agilent Technologies |
| Ec-Phg1.0 | *E. coli* BL21 (DE3) harboring pET-LAAD-HmaS, pRSF-SMDH-AT and pACYC-GluDH | This study |
| Ec-Phg2.0 | *E. coli* BL21 (DE3) harboring pET-LAAD-HmaS and pRSF-SMDH-LeuDH | This study |
| Ec-LAAD | *E. coli* BL21 (DE3) harboring pET-LAAD-HmaS | This study |
